# The T cell receptor repertoire captures healthy aging and CMV independently from epigenetic clocks

**DOI:** 10.64898/2026.02.20.706960

**Authors:** Moira Breëns, Kevin De Man, Yann Heylen, My Ha, Mariia Kuznetsova, Hajar Besbassi, Fabio Affaticati, Wim Vanden Berghe, Benson Ogunjimi, Pieter Meysman

## Abstract

Human aging is the process through which numerous biological changes occur during life, affecting various processes. In some elderly individuals, this functional decline becomes more pronounced, leading to frailty, a condition characterised by reduced physiological reserves and increased vulnerability to stress. With the global rise of life expectancy, identification of biomarkers for healthspan and frailty are becoming more important.

In this study, we directly compare two molecular readouts, namely the T cell receptor (TCR) repertoire and epigenetic clocks, on their ability to discern healthy aging. On blood samples from sixteen individuals, across age-matched healthy elderly and frail individuals, both TCR sequencing and epigenetic profiling were performed. A significantly higher TCR repertoire diversity in the CD4+ T cells differentiated the healthy elderly individuals from the frailty ones. Epigenetic clock signatures of biological relative to chronological ageing rate, did not show a clear difference between both groups. However, when taking into account the CMV-serostatus, a significant increase in epigenetic aging could be observed in the CMV-seropositive individuals.

Our results support a clear hypothesis on the role of CMV infection in the healthy aging of the immune system. In healthy elderly, CMV is typically controlled by CD4+ T cells, however, in the frail elderly, the burden of managing the infection shifts to the CD8+ T cells. This change is marked by two key changes: a decrease in TCR diversity for seropositive individuals compared to seronegative individuals, as well as an increase in the fraction of CMV-associated TCRs within the CD8+ T cells.

These findings contribute to our understanding of aging and provide insight into how CMV-infection may affect healthy aging and frailty. They also underline the crucial role of the immune system in healthy aging and the value of further investigating ageing-related health/disease patterns in the TCR repertoire to determine healthspan/lifespan.

## Introduction

Advances in medicine and technology have led to an increase in the average lifespan and continue to do so. The proportion of individuals older than 60 years is expected to virtually double from 2015 to 2050, from 12% to 22% of the worldwide population. ^1–3^.

Age-related changes are associated with a progressive functional decline, an increased vulnerability to disease, reduced responsiveness to vaccines and an increased risk of age-related diseases.^4^ Healthy aging is the concept of developing and maintaining the functional ability that enables well-being - otherwise referred to as the intrinsic capacity - that enables well-being and independence in older age. ^1–3^ This is captured in the concept of healthspan, the period of good health and performance within one’s life. ^5^

Loss of immune function is one of the main hallmarks of aging, in a process called immunosenescence. The adaptive immune system is severely affected, specifically the T cells, as can be observed by skewing the proportions of CD4+ and CD8+ T cells during the aging process ^6^, as well as the increase in exhausted phenotypes. ^7^ Even the intrinsic diversity of the TCR repertoire is reduced in elderly individuals, which in turn decreased the immune system’s ability to recognize a wide variety of threats. The antigen recognition level, particularly the T cell receptors (TCRs) is also affected by immunosenescence as the TCR repertoire diversity decreases with chronological age. ^8^

Various methods of estimating biological age have been explored in the past, particularly aging clocks - machine learning models that employ biological data to estimate the biological age of an individual. Famously, epigenetic data has been proven to be a good predictor, with examples like the Horvath, Hannum and GrimAge clocks, with the latter integrating phenotypic data with the epigenetic data. The addition of clinical factors also improves disease risk prediction, disease classification and prediction of all-cause mortality. ^9–12^ These epigenetic clocks are also largely influenced by age-related shifts of immune cell composition. More recently, proteomics data and other types of biological data, e.g. immune biomarkers and neuroimaging changes, have been shown to predict biological age, leading to interest in combining multiple levels of biological data in multi-omics aging clocks.^13–16^ It is well known that a chronic cytomegalovirus (CMV) infection is one of the major driving forces of immunosenescence, as it influences age-related changes to several immune parameters. ^17,18^. Additionally, CMV infection has been shown to accelerate epigenetic aging, as higher predicted epigenetic ages were demonstrated in CMV-seropositive individuals compared to their seronegative counterparts. ^19^

Immune cell composition has been shown to affect the outcome of epigenetic aging clocks, as the epigenetic age of T cells is determined by its replicative history, which is driven in turn by cumulative antigen exposure. This is evidenced by the activation of T cells and NK cells being a major driver of epigenetic progression ^20–22^. A single-cell transcriptomics based immune aging clock, as well as a mass cytometry immune clock have been developed recently to capture shifts in e.g. immune cell composition complementary to the pre-existing epigenetic aging clocks. ^23,24^ Additionally, interest in the immune system has risen within the research into frailty, a condition defined by increased vulnerability to stressors, adverse health outcomes and an overall more severe functional decline. Chronic inflammation and immunosenescence are suspected contributors to this condition and they represent potential targets for intervention. ^25^

In this study (Fig. 1), we attempt to unify these distinct observations concerning the dynamics of epigenetic markers and T cell receptor repertoire composition through an in-depth study of a unique age-matched cohort representing healthy aging against frail individuals.

**Figure 1.**
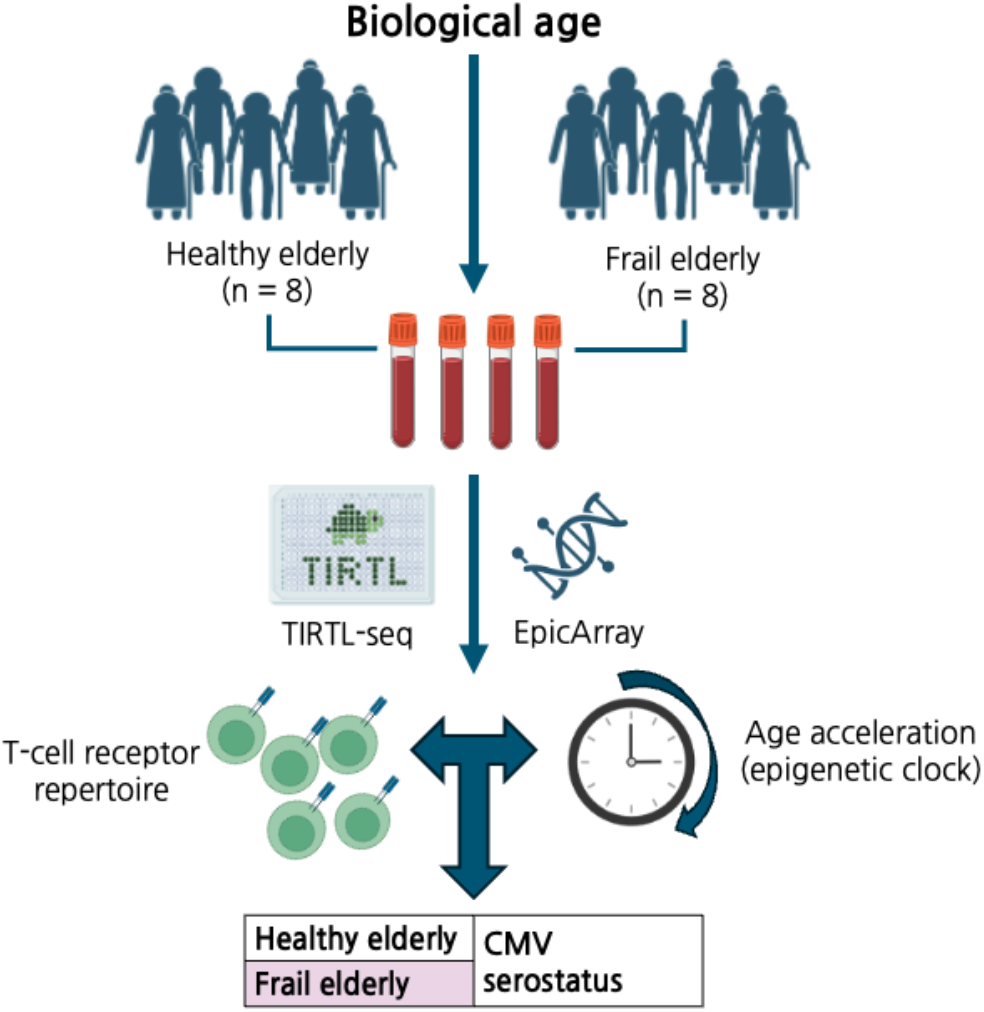
Overview of the study set-up.

## Materials and Methods

### Study cohort

Frail elderly individuals were identified as individuals with a chronological age higher than 65 years old, a moderate to severe vitamin D deficiency (< 20 Vitamin D-OH units/ml) and less than 150 minutes of movement a week (taking both sports and job into account). This refers to the threshold for physical inactivity as set by the WHO. ^26–28^ Healthy elderly individuals are defined as those over 65 years of age, with no vitamin D deficiency and > 150 minutes of movement a week (taking both sports and job into account). Job and time of movement per week were self-reported, and the vitamin D content was measured using the 25-hydroxy Vitamin D test. Cytomegalovirus (CMV) IgG levels were used to determine CMV serostatus, with seronegative being below the detection limit of 0.150 units/ml and CMV seropositive individuals surpassing this limit. Mass cytometry was used to identify the composition of the blood sample on an immune cell level in a subset of the individuals. These measurements are a subset of those reported in Affaticati et al., 2024, and detailed information about the setup in the original publication ^29^.

### TCR sequencing

On blood samples from sixteen age- and sex-matched individuals > 65 years old, equally spread over healthy elderly and frail elderly, both TCR sequencing using TIRTL ^30^ and epigenetic sequencing using EpicArray were performed. The TIRTL-seq protocol was followed as described in Pogorelyy et al., 2025 using RNA extracted from sorted CD4+ and CD8+ T cell samples. One healthy elderly sample failed due to an insufficient number of T cell receptor sequences being detected and difficulties assigning the TCR chain type and was thus excluded from the further analyses. The resulting sequencing reads were processed using the MIXCR package, after which Python was utilised for further processing.

### Epigenetic sequencing

For EpicArray based DNA methylation profiling, peripheral blood mononuclear cells (PBMCs) were cultured for three days, after which they were harvested, counted and pelleted. gDNA was extracted from these pellets using the DNeasy Blood and Tissue Kit (Qiagen, Hilden Germany, 69504) according to manufacturer’s instructions. Subsequently, bisulfite conversion of 250ng isolated DNA was performed using the EZ DNA Methylation Kit (Zymo Research, D5001). Successful conversion was validated by visualization of a methylation-conserved fragment of the human using PyroMark PCR kit (Qiagen, Hilden, Germany, 978703). Amplified products were separated on a 1,5% agarose gel with GelRed™ Nucleic AcidGel staining (Biotium, Fremont, CA, USA, 41002). The TrackIt ™ 100bp DNALadder (Invitrogen; 10488-058) was used as a reference marker. Bisulfite converted samples were hybridised on the Infinium Human Methylation EPIC v2.0 BeadChip (Illumina; 20020531) as described in the manufacturer’s protocol. The EPIC beadchip were analyzed across the genome as previously described ^31,31,32^. Epigenetic clock calculations were performed in R using the methylclock package ^33^ and further processed in a Python environment.

### Statistical testing

Statistical tests such as the independent student’s t-test and the paired student’s t-test were performed using the scipy package, and the plots were built using both matplotlib and seaborn and annotated with Statannotator.

## Results

### Frail aging is typed by vitamin D deficiency and low movement rate

**Table 1.** Overview of the individuals included within the data.

| Features | Frail | Old |
| --- | --- | --- |
| # individuals | 8 | 7 |
| Gender | 50% female | 57% female |
| Age (mean +- standard deviation) | 73 +- 4.4 years | 71 +- 4.7 years |
| BMI (mean +- standard deviation) | 27 +- 3.4 | 27 +- 3.6 |
| Vitamin D (vitamin D-OH units/ml) | 17.1 +- 2.4 | 26.9 +- 7.1 |
| Hours movement/week (mean +- standard deviation) | 1.2 +- 1.3 | 7.7 +- 3.9 |
| CMV serostatus | 62.5% seropositive<br>5/8 individuals | 14.3% seropositive<br>1/7 individuals |
| Availability of Cytof data | 71.4 %<br>4/8 individuals | 57.1 %<br>5/7 individuals |

The final cohort consisted of 15 elderly individuals, of these 15 eight were identified as frail elderly individuals (mean age 72.1 +-4.7 years, 50% female), and the other seven were classified as healthy elderly (mean age 71 +-4.7, 57% female). Healthy elderly individuals are defined as those over 65 years of age, with no vitamin D deficiency (> 40 Vitamin D-OH units/ml) and > 3 hours of movement a week (taking both sports and job into account). Frail elderly individuals were identified as individuals with a chronological age higher than 65 years old, a moderate to severe vitamin D deficiency (< 20 Vitamin D-OH units/ml) and less than three hours of movement a week (taking both sports and job into account). None of the 15 individuals had any comorbidities, nor did any of them smoke.

### Frail aging corresponds to lower CD4+ TCR diversity

Differences in composition of the TCR repertoires between the healthy elderly and frail elderly were examined by using diversity analyses. This repertoire diversity was quantified using Shannon Entropy as a metric, where higher values indicate a more diverse yet evenly distributed set of T cell receptor sequences. Through this metric, we observed no significant difference between the healthy and frail elderly on the combined repertoire level (p-value = 0.0618, Fig. 2A). However, when the TCRs were split by T cell type, namely CD4 and CD8 T cells, a significant decrease in diversity was observed in the CD4+ TCR repertoires in the frail elderly individuals compared to the healthy elderly (p-value = 0.0285). In the CD8+ repertoires no such differences were observed (p-value = 0.97). (Fig. 2B)

**Figure 2.**
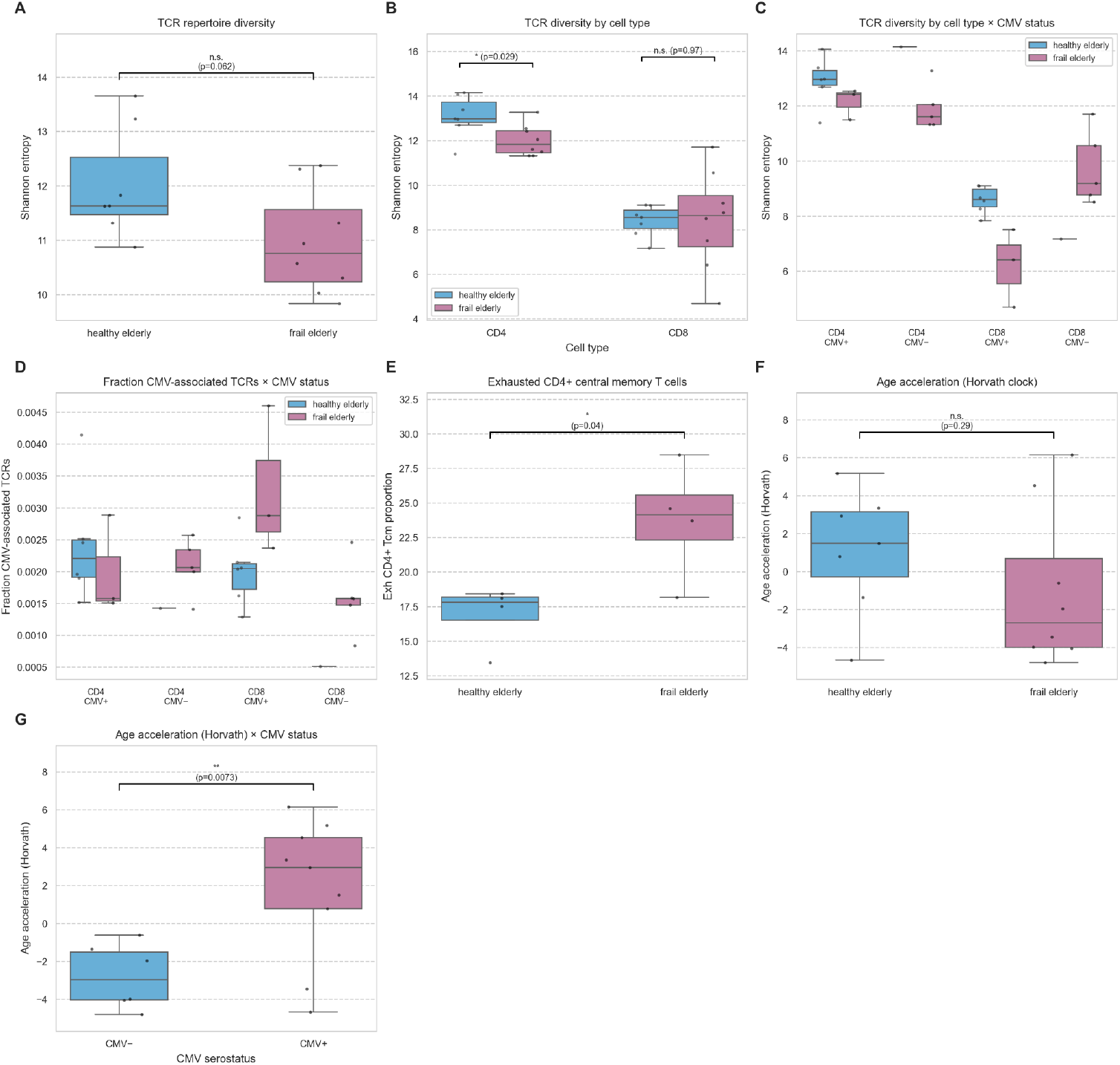
**A)** boxplot displaying TCR repertoire diversity (Shannon entropy) compared between healthy elderly and frail elderly, independent Student’s t-test; **B)** boxplot displaying TCR repertoire diversity (Shannon entropy) compared between healthy elderly and frail elderly within CD4+ T cells and CD8+ T cells, independent Student’s t-test; **C)** boxplot displaying TCR repertoire diversity (Shannon entropy) for each T cell type, distributions are separated by healthy elderly and frail elderly, as well as CMV-serostatus; **D)** boxplot displaying the fraction of CMV-associated TCRs compared between healthy elderly and frail elderly and between CMV serostatus within CD4+ T cells and CD8+ T cells; **E)** boxplot showing a comparison of the fraction of exhausted CD4+ central memory T cells (Cytof) between healthy elderly and frail elderly, independent student t-test; **F)** boxplot showing a comparison between healthy elderly and frail elderly of the Horvath age acceleration 2 (independent student t-test); **G)** boxplot comparing the Horvath age acceleration 2 between CMV serostatus (independent Student’s t-test). Median values are indicated by the horizontal lines within each box, with boxes representing the interquartile range and whiskers extending to 1.5x the interquartile range. Asterisks indicate statistical significance of the independent Student’s t-tests (** p < 0.01, *p < 0.05, n.s. when not significant).

This decrease in TCR repertoire diversity within CD4+ T cells was confirmed in the wider CELLULO EPI dataset, as can be observed in supplementary figure 1. The difference isn’t significant due to the imbalance between healthy elderly (21 individuals) and frail elderly (4 individuals); however, the same trend can be seen. Within the CD8+ T cells no such trends were observed in this dataset once again.

### CMV serostatus skews the CD8+ TCR diversity in frail elderly

As cytomegalovirus (CMV) infections are known to have an effect on immunosenescence and how the immune system ages, it was investigated whether this also impacted the repertoire diversity difference. In the fig. 2C, it can be observed that the CMV serostatus does not affect the Shannon entropy of the CD4+ TCR repertoire and thus does not correlate with the significant difference observed between healthy and frail. In contrast, the CMV serostatus stratified the CD8+ TCR repertoire diversity in the frail elderly individuals, the seropositive frail elderly have a lower Shannon entropy, and the seronegative have a higher Shannon entropy. This was only observed in the frail elderly; no such trend was seen in the healthy elderly. No statistical tests could be performed due to the small sample sizes of the groups.

As CMV serostatus showed differences between the TCR repertoire diversity of the healthy elderly and frail elderly, the proportion of CMV-associated TCRs present in the TCR repertoires was quantified, utilising a list of significantly CMV-associated TCRs (Fig.2E) ^34^. Within the frail elderly, CMV-seropositive individuals show a higher proportion of CMV-associated CD8+ T-cell receptors than the CMV-seronegative individuals. This difference was not observed in the healthy elderly group, however no interpretation can be drawn due to the presence of only a single CMV-seronegative individual. Similar conclusions can be made using ImmuneWatch DETECT, as is demonstrated in supplementary figure 2.^35^

### Frail elderly are typed by an increase of exhausted CD4+ central memory T cells

Comparing the cellular compositions derived from mass cytometry between healthy and frail elderly shows no skewing of the proportions of CD4+ T cells compared to CD8+ T cells (see supplementary figure 3).

As the TCR signature associated with a positive CMV-serostatus is known to be mostly present within the CD4+ memory T cells ^36^, we focused on the CD4+ memory T cells identified using Cytof to compare the healthy elderly and frail elderly. We observed a significant difference in the exhausted CD4+ central memory T cells (Fig. 2D), with these cells being significantly more present in the frail elderly (p-value = 0.031).

### Epigenetic clocks are skewed by CMV serostatus, not by healthy aging

To compare how the immune signals related to epigenetic markers, we applied different Horvath epigenetic clock-based algorithms to estimate biological age. When observing the calculated age acceleration, i.e. the residuals obtained after regressing chronological age and biological age between healthy elderly and frail elderly individuals as predicted by the Horvath epigenetic clock, a trend towards a shallow decrease in this aging parameter can be observed in the frail elderly individuals (Fig.2F).This age acceleration is a second-generation metric for estimating the speed of aging according to the Horvath estimations, with lower values indicating a slower aging process and thus a decrease in biological age.

However as can be observed in supplementary figure 4 the age acceleration is higher in CMV-seropositive individuals than CMV-seronegative individuals, a difference that is observed in both the healthy elderly and frail elderly individuals. When comparing this metric between the CMV-seropositive and CMV-seronegative individuals, it is significantly higher (p-value = 0.0149) in the CMV-seropositive individuals (Fig.2G).

We analysed whether the TCR repertoire data and the epigenetic clock predictions were truly independent of one another (see supplementary figure 5). No relation between the TCR repertoire diversity and the age acceleration was observed, supporting the hypothesis that the adaptive immune system and DNA methylation operate independently, but are both complementarily affected by aging.

## Discussion

Our findings show that frail elderly individuals are typed by a clear, significant decrease in TCR repertoire diversity that could be observed in the CD4+ T cell fraction. This could reflect the compromised state of the immune system that has been associated with frailty and underlines the role of a healthy immune system and a high diversity of the TCR repertoire. This is further supported by the increase of CD4+ exhausted memory T cells found in frail individuals, indicating that a hyperexpansion has occurred.

We observed a difference in TCR repertoire diversity in the CD8+ T cells as well, in this instance related to the CMV serostatus of frail individuals, however statistics were limited due to small sample size. A possible explanation presents itself in the effect of CMV on the immune system. CD4+ T cells play a critical role in the immunological control of viral infections, such as CMV ^37^. If this control is partially lost due to the CD4+ repertoire diversity loss and exhaustion, recruitment of antibodies to manage a CMV infection becomes less feasible. This is supported by the fact that a large fraction of the CMV-associated TCRs are situated in CD4+ memory T cells ^36^. In the absence of a CD4+ response, the immune system may compensate by having CD8+ T cells taking over the immune control role, resulting in a CD8+ inflation and diversity loss in CMV seropositive frail elderly compared to the seronegative frail elderly.

Counterintuitively, the epigenetic clock results show a trend towards a lower biological age in the frail individuals, though not significant, which is opposite from what has been observed previously^38^. However, when stratifying the healthy elderly and frail elderly by their CMV-serostatus, a significant effect could be revealed. In both groups, seropositive individuals have a higher predicted age, and when both elderly groups are combined, this difference is significant. This suggests that the Horvath epigenetic clock reflects the CMV serostatus well, as previous research has indicated, in comparison to the TCR repertoire data which captures an additional, CMV-related difference on top of the difference between healthy aging and frailty. As CMV seroprevalence rises with age ^39,40^, a seropositive status for CMV could be a confounding factor with regards to age, which could explain the inadvertent skewing of these aging clocks towards a CMV-related signal.

It is however important to note that elderly individuals are often underrepresented in epigenetic clock studies which may skew conclusions^41^, an issue that was presented here too, with age underestimations of up to several decades. This could partially be attributed to the difficulties with enrolling (frail) elderly individuals in epigenetic clocks studies, e.g. difficulties in follow-ups leading to missing data, or absence of elderly individuals from the cohort it was trained on ^42^.

Notably, the distinction between healthy elderly and frail elderly was decided by merely two factors, the vitamin D levels and the hours of movement a week. These two factors are modifiable and could be a possible avenue to investigate in the handling of frailty. However, vitamin D is also known to modulate immune functions through the receptors expressed on different immune cell types, such as T cells and B-cells and can as such modulate immune tolerance to a certain extent, as well as playing a crucial role in the development of invariant NK T cells. ^43,44^ It is however remarkable that, based on only two factors, such distinctions are seen on the immune system level. Frailty however is a very ill-defined concept, with a variety of definitions and tests in use ^45^, It is also important to note that this was a retrospective study on an existing cohort, and that the associations seen here do not necessarily indicate a causal relationship, and that the small sample size forms another limitation.

As the immune system is known to be a confounding factor in the training of epigenetic clocks, it was also important to see whether this was affecting our results. An exploratory analysis showed that these were independent from each other. A future avenue of research will be to validate these results in a larger cohort and quantify potential synergies between biomarkers and geroprotective prevention strategies towards healthy aging.

## References

1. Ageing and health. Accessed January 9, 2026. https://www.who.int/news-room/fact-sheets/detail/ageing-and-health

2. Beard JR, Officer A, Carvalho IA de, et al. The World report on ageing and health: a policy framework for healthy ageing. The Lancet. 2016;387(10033):2145–2154. doi:10.1016/S0140-6736(15)00516-4

3. Rudnicka E, Napierała P, Podfigurna A, Męczekalski B, Smolarczyk R, Grymowicz M. The World Health Organization (WHO) approach to healthy ageing. Maturitas. 2020;139:6–11. doi:10.1016/j.maturitas.2020.05.018

4. Pawelec G, Bronikowski A, Cunnane SC, et al. The conundrum of human immune system “senescence.” Mech Ageing Dev. 2020;192:111357. doi:10.1016/j.mad.2020.111357

5. Masfiah S, Kurnialandi A, Meij JJ, Maier AB. Definitions of healthspan: A systematic review. Ageing Res Rev. 2025;111:102806. doi:10.1016/j.arr.2025.102806

6. Han S, Georgiev P, Ringel AE, Sharpe AH, Haigis MC. Age-associated remodeling of T cell immunity and metabolism. Cell Metab. 2023;35(1):36–55. doi:10.1016/j.cmet.2022.11.005

7. Fulop T, Larbi A, Pawelec G. Human T Cell Aging and the Impact of Persistent Viral Infections. Front Immunol. 2013;4. doi:10.3389/fimmu.2013.00271

8. Britanova OV, Putintseva EV, Shugay M, et al. Age-Related Decrease in TCR Repertoire Diversity Measured with Deep and Normalized Sequence Profiling. J Immunol. 2014;192(6):2689–2698. doi:10.4049/jimmunol.1302064

9. Hannum G, Guinney J, Zhao L, et al. Genome-wide Methylation Profiles Reveal Quantitative Views of Human Aging Rates. Mol Cell. 2013;49(2):359–367. doi:10.1016/j.molcel.2012.10.016

10. Horvath S. DNA methylation age of human tissues and cell types. Genome Biol. 2013;14(10):3156. doi:10.1186/gb-2013-1410-r115

11. Lu AT, Quach A, Wilson JG, et al. DNA methylation GrimAge strongly predicts lifespan and healthspan. Aging. 2019;11(2):303–327. doi:10.18632/aging.101684

12. Mavrommatis C, Belsky DW, Ying K, et al. An unbiased comparison of 14 epigenetic clocks in relation to 10-year onset of 174 disease outcomes in 18,859 individuals. medRxiv. Preprint posted online July 15, 2025:2025.07.14.25331494. doi:10.1101/2025.07.14.25331494

13. Argentieri MA, Xiao S, Bennett D, et al. Proteomic aging clock predicts mortality and risk of common age-related diseases in diverse populations. Nat Med. 2024;30(9):2450–2460. doi:10.1038/s41591-024-03164-7

14. Guo X, Robertson JA, Aparicio A, et al. Variations in Innate Immune Cell Subtypes Correlate with Epigenetic Clocks, Inflammaging and Health Outcomes. Adv Sci. 2025;12(43):e05922. doi:10.1002/advs.202505922

15. Min M, Egli C, Dulai AS, Sivamani RK. Critical review of aging clocks and factors that may influence the pace of aging. Front Aging. 2024;5. doi:10.3389/fragi.2024.1487260

16. Wen J. Refining the generation, interpretation and application of multi-organ, multi-omics biological aging clocks. Nat Aging. 2025;5(9):1897–1913. doi:10.1038/s43587-025-00928-9

17. Pawelec G, McElhaney JE, Aiello AE, Derhovanessian E. The impact of CMV infection on survival in older humans. Curr Opin Immunol. 2012;24(4):507–511. doi:10.1016/j.coi.2012.04.002

18. Pawelec G, Derhovanessian E. Role of CMV in immune senescence. Virus Res. 2011;157(2):175–179. doi:10.1016/j.virusres.2010.09.010

19. Kananen L, Nevalainen T, Jylhävä J, et al. Cytomegalovirus infection accelerates epigenetic aging. Exp Gerontol. 2015;72:227–229. doi:10.1016/j.exger.2015.10.008

20. Jonkman TH, Dekkers KF, Slieker RC, et al. Functional genomics analysis identifies T and NK cell activation as a driver of epigenetic clock progression. Genome Biol. 2022;23(1):24. doi:10.1186/s13059-021-02585-8

21. Mi T, Soerens AG, Alli S, et al. Conserved epigenetic hallmarks of T cell aging during immunity and malignancy. Nat Aging. 2024;4(8):1053–1063. doi:10.1038/s43587-024-00649-5

22. Tomusiak A, Floro A, Tiwari R, et al. Development of an epigenetic clock resistant to changes in immune cell composition. Commun Biol. 2024;7(1):934. doi:10.1038/s42003-024-06609-4

23. Li W, Zhang Z, Kumar S, et al. Single-cell immune aging clocks reveal inter-individual heterogeneity during infection and vaccination. Nat Aging. 2025;5(4):607–621. doi:10.1038/s43587-025-00819-z

24. Sim HB, Jang JH, Mun SK, et al. Reading the immune clock: a machine learning model predicts mouse immune age from cellular patterns. Nat Commun. 2025;17(1):640. doi:10.1038/s41467-025-67393-1

25. Pilotto A, Custodero C, Maggi S, Polidori MC, Veronese N, Ferrucci L. A multidimensional approach to frailty in older people. Ageing Res Rev. 2020;60:101047. doi:10.1016/j.arr.2020.101047

26. Marcos-Pérez D, Sánchez-Flores M, Proietti S, et al. Low Vitamin D Levels and Frailty Status in Older Adults: A Systematic Review and Meta-Analysis. Nutrients. 2020;12(8):2286. doi:10.3390/nu12082286

27. Wang X, Liang Y, Jin C, et al. Combined effects of vitamin D and cumulative dietary risk score on fatty liver and mortality in vulnerable individuals: a prospective analysis from the UK Biobank. GeroScience. Published online July 1, 2025. doi:10.1007/s11357-025-01762-y

28. WHO guidelines on physical activity and sedentary behaviour. Accessed January 11, 2026. https://iris.who.int/items/2abc9ed0-2a55-4a07-a65f-ab4b7c412ecc

29. Affaticati F, Ha MK, Gehrmann T, et al. Bridging immunotypes and enterotypes using a systems immunology approach. bioRxiv. Preprint posted online November 29, 2024:2024.11.29.625344. doi:10.1101/2024.11.29.625344

30. Pogorelyy MV, Kirk AM, Adhikari S, et al. TIRTL-seq: deep, quantitative and affordable paired TCR repertoire sequencing. Nat Methods. 2026;23(1):56–64. doi:10.1038/s41592-025-02907-9

31. D’Incal C, Van Dijck A, Ibrahim J, et al. ADNP dysregulates methylation and mitochondrial gene expression in the cerebellum of a Helsmoortel–Van der Aa syndrome autopsy case. Acta Neuropathol Commun. 2024;12(1):62. doi:10.1186/s40478-024-01743-w

32. Logie E, Van Puyvelde B, Cuypers B, et al. Ferroptosis Induction in Multiple Myeloma Cells Triggers DNA Methylation and Histone Modification Changes Associated with Cellular Senescence. Int J Mol Sci. 2021;22(22):12234. doi:10.3390/ijms222212234

33. Pelegí-Sisó D, de Prado P, Ronkainen J, Bustamante M, González JR. methylclock: a Bioconductor package to estimate DNA methylation age. Bioinformatics. 2021;37(12):1759–1760. doi:10.1093/bioinformatics/btaa825

34. Emerson RO, DeWitt WS, Vignali M, et al. Immunosequencing identifies signatures of cytomegalovirus exposure history and HLA-mediated effects on the T cell repertoire. Nat Genet. 2017;49(5):659–665. doi:10.1038/ng.3822

35. ImmuneWatch DETECT | IMW-DETECT. Accessed February 19, 2026. https://your-docusaurus-site.example.com/detect-docs/

36. De Neuter N, Bartholomeus E, Elias G, et al. Memory CD4+ T cell receptor repertoire data mining as a tool for identifying cytomegalovirus serostatus. Genes Immun. 2019;20(3):255–260. doi:10.1038/s41435-018-0035-y

37. Gamadia LE, Remmerswaal EBM, Weel JF, Bemelman F, van Lier RAW, Ten Berge IJM. Primary immune responses to human CMV: a critical role for IFN-γ–producing CD4+ T cells in protection against CMV disease. Blood. 2003;101(7):2686–2692. doi:10.1182/blood-2002-08-2502

38. You Y, Chen Y, Ding H, et al. Relationship between physical activity and DNA methylation-predicted epigenetic clocks. Npj Aging. 2025;11(1):27. doi:10.1038/s41514-025-00217-0

39. Fowler K, Mucha J, Neumann M, et al. A systematic literature review of the global seroprevalence of cytomegalovirus: possible implications for treatment, screening, and vaccine development. BMC Public Health. 2022;22(1):1659. doi:10.1186/s12889-022-13971-7

40. Lachmann R, Loenenbach A, Waterboer T, et al. Cytomegalovirus (CMV) seroprevalence in the adult population of Germany. PLoS ONE. 2018;13(7):e0200267. doi:10.1371/journal.pone.0200267

41. El Khoury LY, Gorrie-Stone T, Smart M, et al. Systematic underestimation of the epigenetic clock and age acceleration in older subjects. Genome Biol. 2019;20(1):283. doi:10.1186/s13059-019-1810-4

42. Föhr T, Törmäkangas T, Lankila H, et al. The Association Between Epigenetic Clocks and Physical Functioning in Older Women: A 3-Year Follow-up. J Gerontol Ser A. 2022;77(8):1569–1576. doi:10.1093/gerona/glab270

43. Aranow C. Vitamin D and the Immune System. J Investig Med. 2011;59(6):881–886. doi:10.2310/JIM.0b013e31821b8755

44. Cantorna MT, Zhao J, Yang L. Vitamin D, invariant natural killer T-cells and experimental autoimmune disease. Proc Nutr Soc. 2012;71(1):62–66. doi:10.1017/S0029665111003193

45. Kudelka J, Ollenschläger M, Dodel R, et al. Which Comprehensive Geriatric Assessment (CGA) instruments are currently used in Germany: a survey. BMC Geriatr. 2024;24(1):347. doi:10.1186/s12877-024-04913-6

