## Supplementary figures for "The T cell receptor repertoire captures healthy aging and CMV independently from epigenetic clocks"

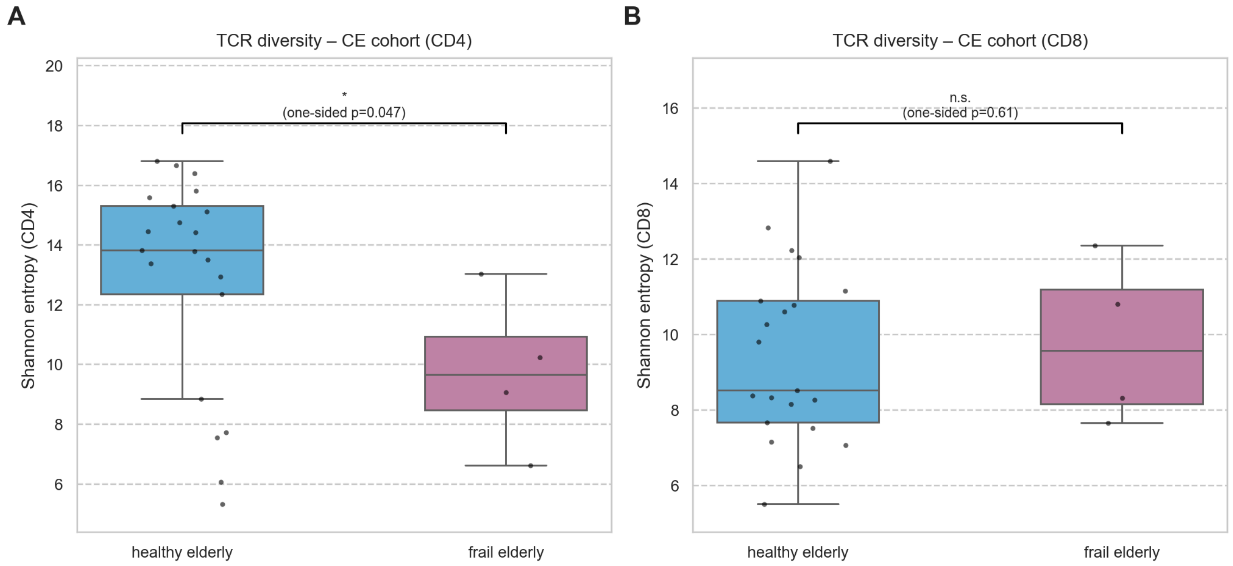


**Supplementary figure 1: A)** boxplot displaying the TCR repertoire diversity (Shannon entropy) compared between healthy elderly and frail elderly in the CD4+ T cell fraction, one-sided Welch's t-tests to confirm the observed decrease in TCR repertoire diversity in the frail elderly CD4+ fraction ; **B)** boxplot displaying the TCR repertoire diversity (Shannon entropy) compared between healthy elderly and frail elderly in the CD8+ T cell fraction, independent Student's t-test;
Median values are indicated by the horizontal lines within each box, with boxes representing the interquartile range and whiskers extending to 1.5x the interquartile range.
Asterisks indicate statistical significance of the independent Student's t-tests (** $p < 0.01$, *$p < 0.05$, n.s. when not significant).

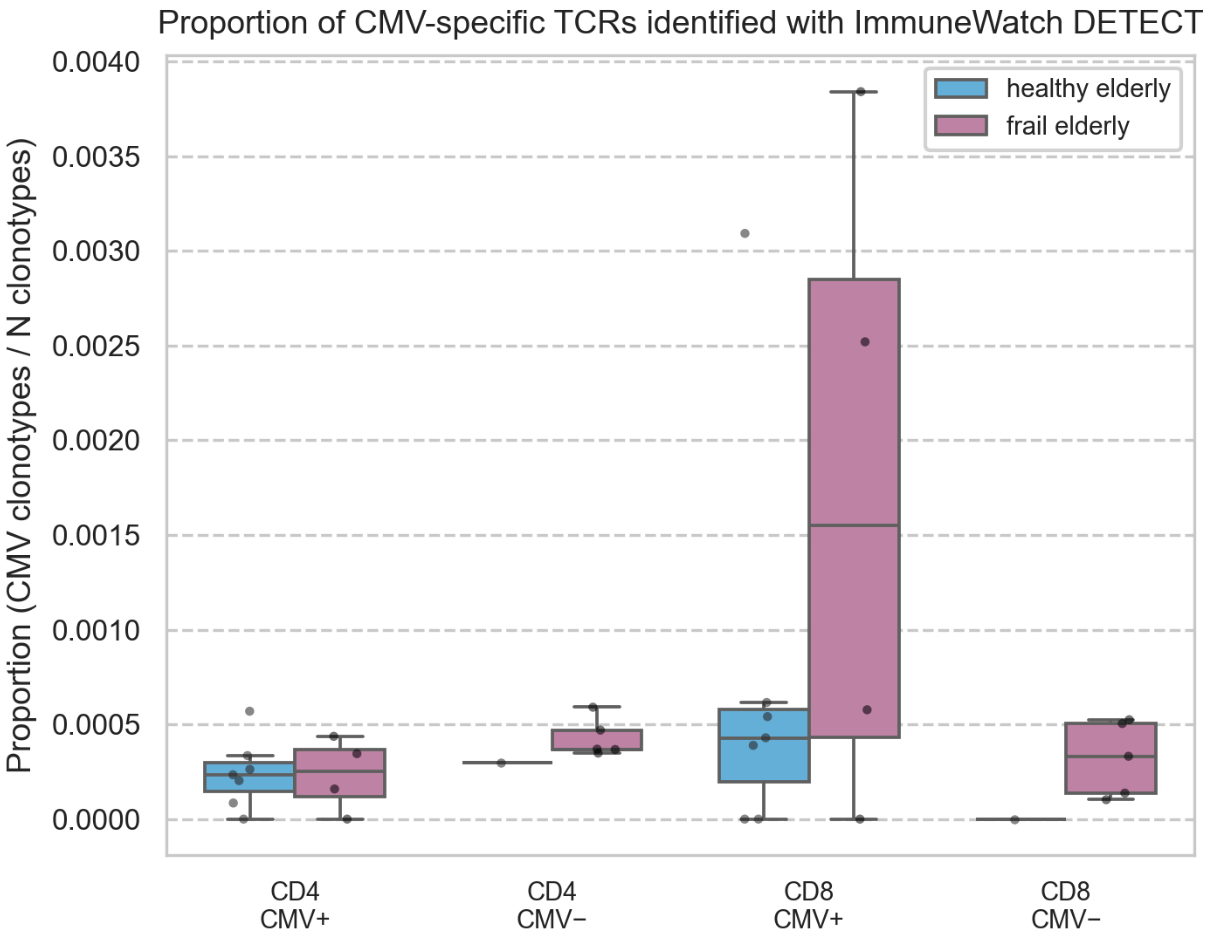


**Supplementary figure 2:** a boxplot displaying the fraction of CMV-associated TCRs compared between healthy elderly and frail elderly and between CMV serostatus within CD4+ T cells and CD8+ T cells; the recommended cut-off of 0.23 was used for the ImmuneWatch DETECT score to filter the results
Median values are indicated by the horizontal lines within each box, with boxes representing the interquartile range and whiskers extending to 1.5x the interquartile range.
Asterisks indicate statistical significance of the independent Student's t-tests (** $p < 0.01$, *$p < 0.05$, n.s. when not significant).

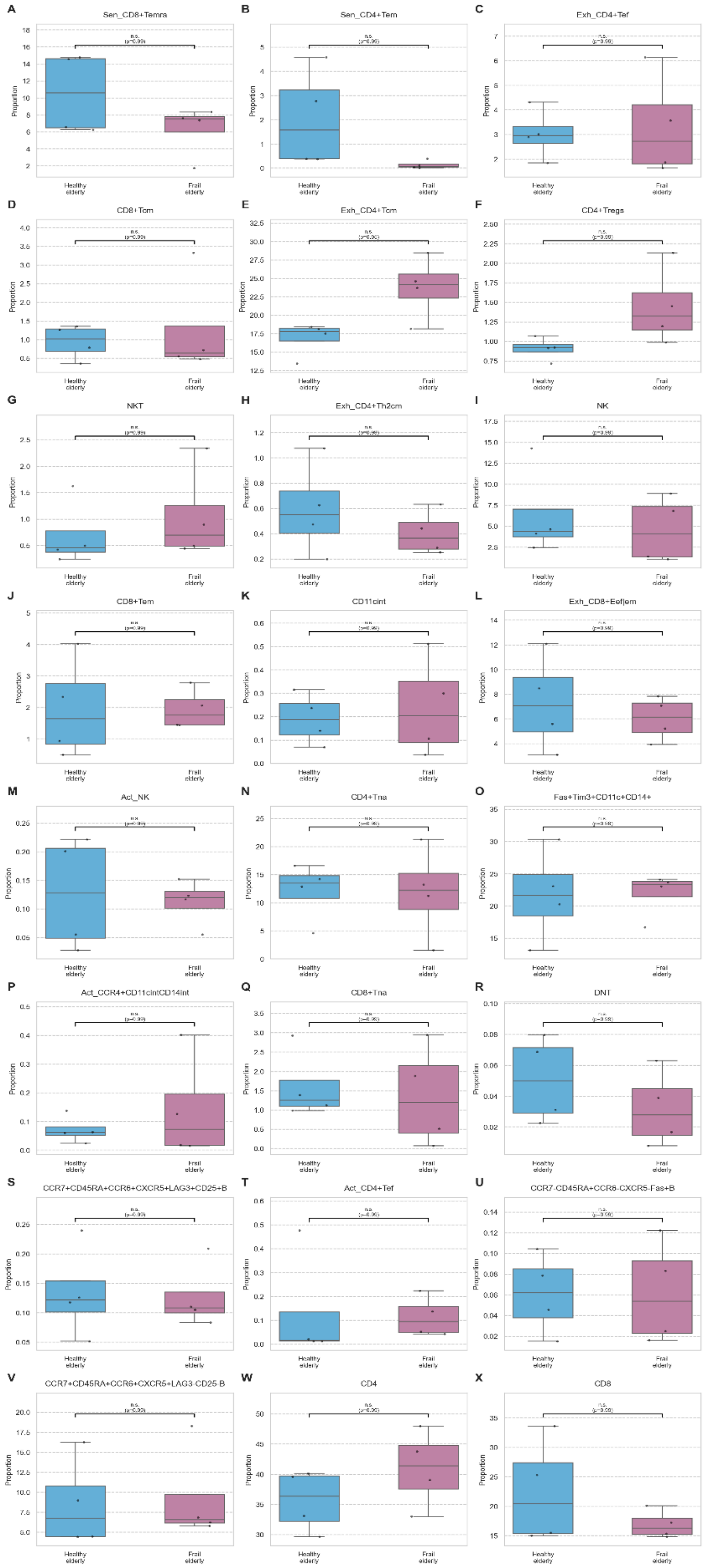
A
Sen_CD8+Temra
B
Sen_CD4+Tem
C
Exh_CD4+Tef
D
CD8+Tcm
E
Exh_CD4+Tcm
F
CD4+Tregs
G
NKT
H
Exh_CD4+Th2cm
I
NK
J
CD8+Tem
K
CD11cint
L
Exh_CD8+Eef|em
M
Act_NK
N
CD4+Tna
O
Fas+Tim3+CD11c+CD14+
P
Act_CCR4+CD11cintCD14int
Q
CD8+Tna
R
DNT
S
CCR7+CD45RA+CCR6+CXCR5+LAG3+CD25+B
T
Act_CD4+Tef
U
CCR7-CD45RA+CCR6-CXCR5-Fas+B
V
CCR7+CD45RA+CCR6+CXCR5+LAG3-CD25-B
W
CD4
X
CD8
Proportion
Healthy elderly
Frail elderly
n.s.

**Supplementary figure 3:** A boxplot showing comparisons of all the fractions observed using Cytof between healthy elderly and frail elderly, as well as comparisons for the entire CD4+ and the entire CD8+ fraction; independent student t-test with Benjamini-Hochberg FDR multiple testing correction

Median values are indicated by the horizontal lines within each box, with boxes representing the interquartile range and whiskers extending to 1.5x the interquartile range.

Asterisks indicate statistical significance of the independent Student's t-tests (** $p < 0.01$, *$p < 0.05$, n.s. when not significant).

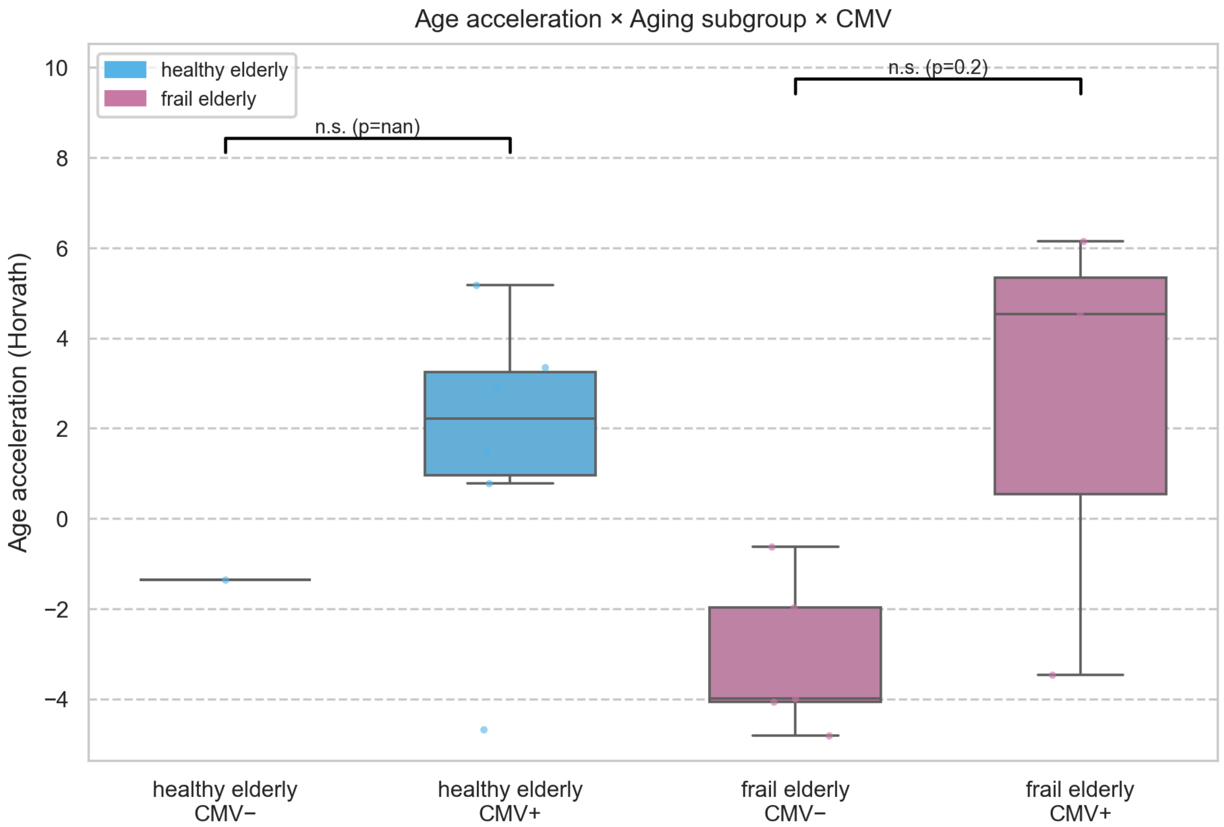


**Supplementary figure 4:** a boxplot comparing age acceleration of the Horvath epigenetic clock, between CMV serostatus within the healthy elderly and frail elderly, independent Student t-test

Median values are indicated by the horizontal lines within each box, with boxes representing the interquartile range and whiskers extending to 1.5x the interquartile range.

Asterisks indicate statistical significance of the independent Student's t-tests (** $p < 0.01$, *$p < 0.05$, n.s. when not significant).

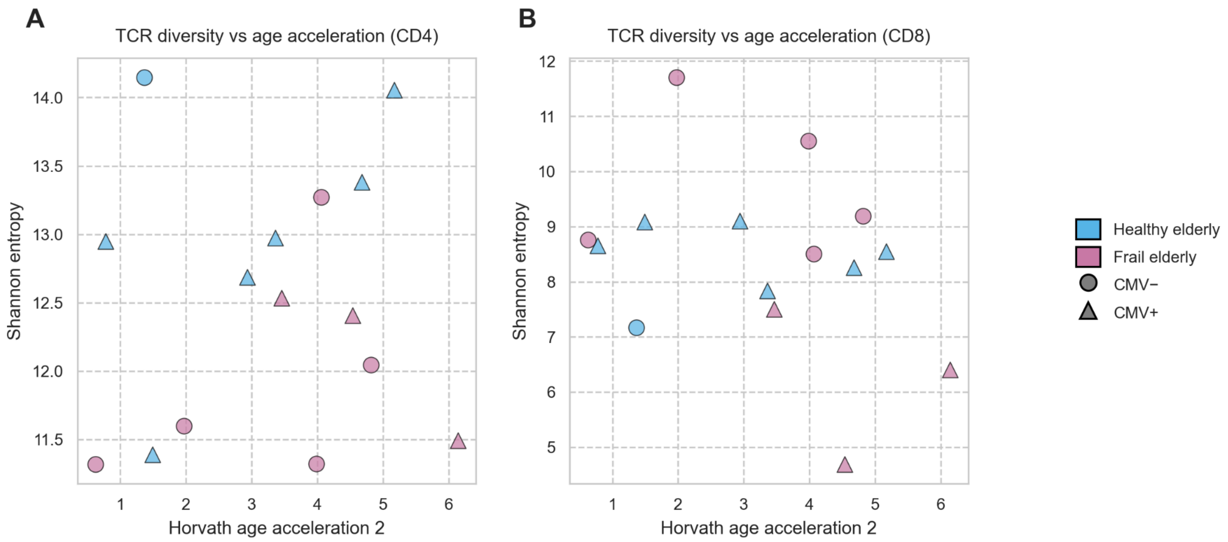


**Supplementary figure 5: A)** Scatterplot comparing the Horvath age acceleration and the TCR repertoire diversity (Shannon entropy) between healthy and frail elderly, further split up by CMV serostatus, in the CD4+ fraction; **B)** Scatterplot comparing the Horvath age acceleration and the TCR repertoire diversity (Shannon entropy) between healthy and frail elderly, further split up by CMV serostatus, in the CD8+ fraction; legend for both scatterplots is depicted on the far right
